# Evaluation of the impact of *Shigella* virulence genes on the basis of clinical features observed in patients with shigellosis

**DOI:** 10.1101/865832

**Authors:** Visnu Pritom Chowdhury, Ishrat Jahan Azmi, Mohammad Rafiqul Islam, Mahmuda Akter, Shahin Mahmud, Abu Syed Golam Faruque, Kaisar Ali Talukder

**Affiliations:** Biotechnology and Microbiology Program, Department of Mathematics and Natural Sciences, BRAC University, Dhaka, Bangladesh; Enteric and Food Microbiology Laboratory, Laboratory Sciences Division, icddr,b, Dhaka, Bangladesh; Nutrition and Clinical Services Division, icddr,b, Dhaka, Bangladesh; Department of Biotechnology and Genetic Engineering, Mawlana Bhashani Science and Technology University, Tangail, Bangladesh

**Keywords:** shigellosis, diarrhoea, *Shigella*, T3SS, enterotoxin

## Abstract

*Shigella* is still attributable to nearly 164,300 deaths annually, mostly in sub-Saharan Africa and South Asian young children, remaining as a major public health threat, especially in the developing countries. Our goal was to study the association between *Shigella* virulence genes and clinical features observed in shigellosis. Therefore, 61 *S. flexneri* strains were investigated, isolated from patients from a tertiary level facility in Bangladesh between 2009 to 2013. Subsequently, the presence of 140 MDa large virulence plasmid (p140), virulence (*ipaH, ial*), toxin (*set, sen*) and T3SS related genes (*virB, ipaBCD, ipgC, ipgB1, ipgA, icsB, ipgD, ipgE, ipgF, mxiH, mxiI, mxiK, mxiE, mxiC, spa15, spa47, spa32* and *spa24*) were evaluated. p140 was found in 79% (n=48) cases. *ipaBCD* was found in 90% (n=55) strains, while seven of them were missing p140. However, *ial* was found in 89% isolates, and *ipgC* and *ipgE* in 85% cases. The prevalence of the rest of the genes was less than 85%. These findings were then compared against the clinical features of each of the corresponding pathogens, and several statistically significant correlations were observed (all p<0.05). Briefly, the enterotoxin genes (*set, sen*) and another virulence gene (*ial*) were found significantly associated with several clinical features of shigellosis, including bloody mucoid stool, rectal straining, fever, and abdominal pain. Our findings reiterate that the diarrheal disease severity is significantly associated with the enterotoxin producing *Shigella* infection, also suggesting that the T3SS related virulence genes might be translocated elsewhere other than the 140 MDa large virulence plasmid.

## Introduction

Even though the incidence of shigellosis has been reduced significantly worldwide through improved sanitation and safer water supply, it is still one of the major causes of fatalities from diarrhoeal diseases annually, especially in sub-Saharan Africa and South Asian under-five children (1, 2)□. There are four members of this species - *S. dysenteriae, S. flexneri, S. boydii*, and *S. sonnei*, and *Shigella dysenteriae* type 1 is responsible for the most severe form of shigellosis. However, no incidence of *S. dysenteriae* type 1 has been reported since 2004. For the last 35 years, *S. flexneri* has been found as the predominant species in Bangladesh (3)□, while we observed a major swing in the epidemiology of *Shigella* species – mostly over the last two decades. In 2010, the prevalence of *S. sonnei* was 27%, while in 2000 it was 8% (3, 4)□. *Shigella* can be easily transmittable through the fecal-oral route. With no effective vaccine, low infectious dose - approximately 10-100 organisms (5)□ - and with the emergence of multi-drug resistant *Shigella* spp. (6)□, it is still wise to consider *Shigella* as a major threat, that can wreak havoc on any public health system in any developing tropical country, especially one like Bangladesh.

After getting into the gastrointestinal tract these pathogens pass through the acid barrier of the stomach, multiply in the intestine and reach the colon, where they invade colonic mucosa and multiply again. Afterward, these organisms are taken up by the macrophages, where they induce cell death, causing the release of proinflammatory cytokines outside, attracting the natural killer cells and the polymorphonuclear neutrophils. These events destroy the gut mucosa cell lining and result in the dysenteric illness (7)□.

The cellular pathogenesis and clinical presentation of dysenteric illness are mediated by multiple *Shigella* virulence factors including, p140, enterotoxins and Type 3 Secretion system (T3SS). The p140 is a 200 kb long large virulence plasmid containing 100 genes with a 31 kb conserved “entry region”. This part of the plasmid is composed of 34 genes, divided into 4 functional groups. Group I contains the effector genes, encoding the invasion plasmid antigen (Ipa), secreted via the T3SS. These Ipa proteins manipulate the host cell processes in favor of the pathogen (8)□. Group II consists of membrane expression of ipa (*mxi*) and surface presentation of *ipa* (*spa*) genes – encoding Mxi and Spa proteins - required to secrete the Ipa and other effector proteins (9)□. Group III is made of two regulatory genes-*virB* and *mxiE* - regulating the T3SS genes (10)□. And Group IV contains chaperone protein-coding genes, required to stabilize the T3SS substrates within the bacterial cytoplasm (11)□. However, all operations of p140 are strictly controlled by the global regulatory elements described elsewhere (7)□.

The toxins are an essential virulence factor for *Shigella* species. And unlike ShET2 - encoded by the *sen* gene, located in p140 - Shiga toxin (Stx) and *Shigella* enterotoxin 1 (ShET1) are encoded in the chromosome. Stx is found in *S. dysenteriae* type 1 - also in enterohemorrhagic *E. coli* (12)□ - and ShET1 is seen in *S. flexneri* (13)□. However, Stx is responsible for a more severe form of complications, including hemolytic-uremic syndrome, even death (14)□.

The T3SS is a needle-like apparatus, composed of a base, needle, inner membrane export apparatus, cytosolic components, tip complex, and the translocons. The base spans over both outer membranes, composed of 2 concentric rings. The outer ring is formed with MxiD protein, and the inner ring is made of MxiG and MxiJ proteins (15)□. The outer rod of the needle is made of mostly by multiple (∼100) copies of MxiH protein, arranged in a helical pattern. However, the inner rod is made of MxiI, determining the needle-length. The inner membrane export apparatus consists of five conserved proteins - MxiA, Spa24, Spa9, Spa29, Spa40 - acting as a protein channel and facilitating the target proteins through the inner membrane (15)□. The cytosolic components recruit, unfold and transport the substrates following contact with the host cell, while a linker protein - MxiN - recruits the cytosolic components - e.g., ATPase - to the sorting platform (16)□. The activation of the needle complex occurs following the contact with the host cell, mediated by the tip complex - i.e., IpaB, IpaC, IpaD (17)□ - leading toward the passage of effectors into the host cell and initiating the pathogenesis. These cellular events often lead to serious illness in humans, ranging from diarrhea with tenesmus and high fever to serious life-threatening complications such as Shigella encephalopathy, hemolytic uremic syndrome, even death.

Given the fact that shigellosis is a major public health concern in Bangladesh and its molecular background is well known, this study is aimed to evaluate the relationship between *Shigella* virulence (*ipaH, ial*), toxin (*set, sen*), and T3SS related genes (*virB, ipaBCD, ipgC, ipgB1, ipgA, icsB, ipgD, ipgE, ipgF, mxiH, mxiI, mxiK, mxiE, mxiC, spa15, spa47, spa32, spa24*) and the clinical features observed in patients with shigellosis.

## Materials and methods

### Bacterial isolates

Sixty-one clinical isolates of *S. flexneri* of different serotypes were obtained from the patients enrolled in “Disease burden and etiologic agents of diarrhea patients visiting Kumudini Hospital, Mirzapur” study from 2009 to 2013. In the 61 *S. flexneri* strains serotype 1b, 1c, 2a, 2b, 3a, 4, and 6 were 5, 9, 16, 15, 8, 3, and 5 respectively. All these strains were isolated and characterized at the Enteric and Food Microbiology Laboratory of icddr,b following standard Microbiological and Biochemical methods (18)□. A single colony of confirmed *Shigella* strain was grown in Tryptic Soy Broth (TSB) with 0.3% yeast extract and stored at −70 °C after adding 15% glycerol for further use. For plasmid analysis and PCR assay, *S. flexneri* 2a strain YSH6000 (19)□ and *Escherichia coli* strain ATCC 25922 were used as the positive and negative control, respectively. The following *E. coli* strains PDK-9, V-517, Sa, and R1 were used as plasmid molecular weight standards. Strains that were used as standards were collected from the Enteric and Food Microbiology Laboratory of icddr,b. Information on clinical features corresponding to the 61 samples was extracted from the database.

### Serological Typing

Each strain was serologically confirmed using a commercially available antisera kit - Denka Seiken, Co. Ltd. Japan - as well as a monoclonal antibody - Reagensia AB, Stockholm, Sweden - reagents, specific for all *S. flexneri* type and group-factor antigens. Strains were subcultured on MacConkey agar-Difco, Becton Dickinson & Company Sparks, MD, USA - plates and after overnight incubation serological reactions were carried out by the glass slide agglutination test as described previously by El-Gendy et al., 1999 (18)□.

### Plasmid analysis

Plasmid DNA was processed according to the alkaline lysis method of Kado and Liu (20)□ with few modifications (21)□. An isolated colony for each strain was inoculated in 1.5 mL TSBY - Tryptic Soy broth with 0.3% yeast extract - and incubated overnight at 37 °C on a water-bath shaker. Later the cells were harvested by centrifugation and were suspended in 100 μL of solution I (40 mM Tris-Na acetate, 2 mM EDTA, pH 7.4). Then 200 μL of solution II - 3% sodium dodecyl sulfate, 50 mM Tris, pH 12.9 - was added and incubated at 55 °C for 1 hour. After incubation, an equal volume of solution III -phenol-chloroform-isoamyl alcohol [25:24:1] - was added and mixed carefully before collecting the plasmid DNA by centrifugation. Then the plasmid DNA was separated by horizontal gel electrophoresis in a 0.7% agarose gel at room temperature at 100V (30 mA) in TBE (Tris-borate-EDTA) buffer, taking approximately 3 hours. The slab gel was stained with ethidium bromide and the DNA band images were taken by a gel documentation system. The masses of unknown plasmid DNA were measured against comparing the mobilities with the known molecular weights (22)□. Plasmids present in strains *E. coli* PDK-9 (140, 105, 2.7 and 2.1 MDa), V-517 (35.8, 4.8, 3.7, 3.4, 3.1, 2.0, 1.8 and 1.4 MDa), Sa (23 MDa) and R1 (62 MDa) were used as molecular weight standards.

### PCR assay

The detection of the following genes in *Shigella* isolates was carried out by PCR assay: *ipaH, ial, set* (*Shigella* enterotoxin 1), *sen* (*Shigella* enterotoxin 2) and T3SS related genes - *virB, ipaBCD, ipgC, ipgB1, ipgA, icsB, ipgD, ipgE, ipgF, mxiH, mxiI, mxiK, mxiE, mxiC, spa15, spa47, spa32* and *spa24* - according to the procedure of Vargas, et al., 1999 (23)□. However, PCR protocols for *spa15* and *mxiE* were developed in this study after designing the respective PCR primers, following instructions described elsewhere (24)□. During PCR, *S. flexneri* 2a strain YSH6000 was used as the positive control, and *E. coli* ATCC 25922 lacking the p140 was used as the negative control.

### Statistical analysis

Cross tabulations followed by Chi-square tests of independence were carried out to test the null hypothesis, whether the presence of a particular clinical feature does not depend upon the presence of a specific gene. The results were considered statistically significant if the p-value was found to be <0.05. All statistical analyses were carried out with IBM SPSS Statistics Version 22.0 software. Graphs and figures were prepared using Image Lab Version 5.2.1, IBM SPSS Statistics Version 22.0, Image J 1.48k, GNU Image Manipulation Program Version 2.8.10, and draw.io Version 11.3.0 software.

## Results

### Serological typing

All 61 isolates were *S. flexneri*, where serotype 1b, 1c, 2a, 2b, 3a, 4, and 6a were, 5, 9, 16, 15, 8, 3, and 5 respectively.

### Plasmid profile analysis

Of the 61 isolates, 79% (48/61) were p140 positive. Among these 48 strains, 15, 14, 8, 3, and 5 were, *S. flexneri* 2a, 2b, 3a, Type-4, and 6a, respectively. None of the 1b strains was found p140 positive, while, three 1c strains (3/9) were found positive for p140. The small plasmids - approximately 2.7 and 2.1 MDa - were observed in around 94% strains, thus regarded as core plasmids. Nonetheless, the overall plasmid population was heterogeneous (Fig 1).

**Fig 1.**
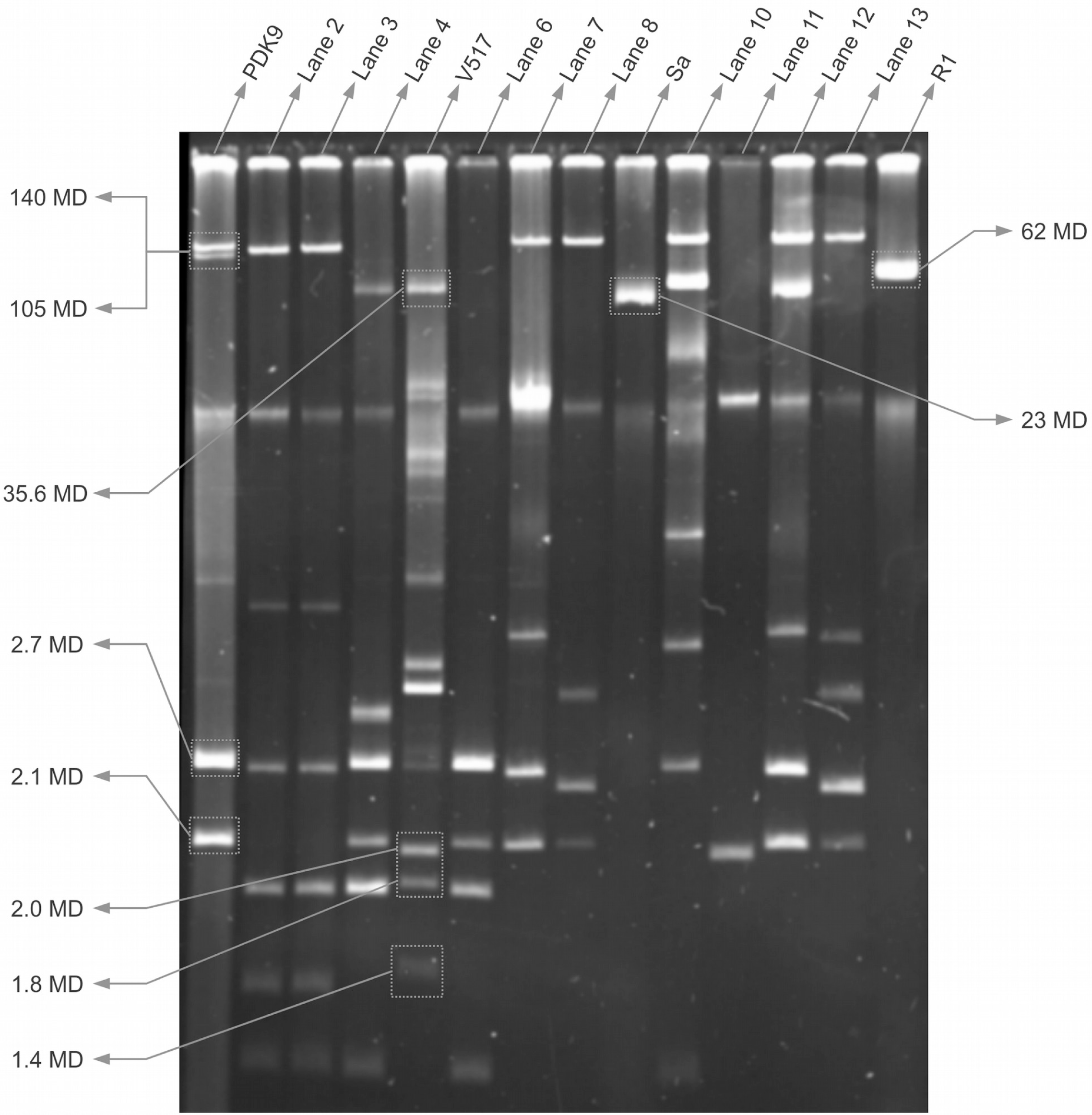
Plasmid DNA analysis of representative *S. flexneri* strains. Gel electrophoresis of characterized plasmid DNA. Lane 1, *E. coli* PDK-9; Lane 2, K-4081 *S. flexneri* 6a; Lane 3, K-2336 *S. flexneri* 6a; Lane 4, K-3253 *S. flexneri* 1c; Lane 5, *E. coli* V-517; Lane 6, K-3127 *S. flexneri* 1b; Lane 7, K-2980 *S. flexneri* 2a; Lane 8, K-1502 *S. flexneri* 3a; Lane 9, *E. coli* Sa; Lane 10, K-632 *S. flexneri* 4; Lane 11, K-553 *S. flexneri* 4; Lane 12, K-102 *S. flexneri* 2b; Lane 13, K-6 *S. flexneri* 3b; Lane 14, *E. coli* R1. The reference position of the reference strains at 140 MDa, 105 MDa, 62 MDa, 35.6 MDa, 23 MDa, 2.7 MDa, 2.1 MDa, 2.0 MDa, 1.8 MDa and 1.4 MDa plasmid DNA shown by indicators.

### Assessment of enterotoxin and virulence genes

All of the 61 *S. flexneri* strains were analyzed to evaluate the presence of virulence (*ipaH, ial*) and toxin (*set, sen*) genes by PCR assay. All were found *ipaH* positive, while *ial* was found in 54 isolates. All of the 7 *ial* negative strains were also p140 negative, but 6 *ial* positive strains were found p140 negative. *sen* was found positive in 49 strains and was found negative in 12. All of the *sen* negative strains were also p140 negative. In the case of *set*, 34 *S. flexneri* strains were found *set* positive and 27 were *set* negative. Out of the 27 *set* negative strains, 22 were both *set1A* and *set1B* negative, and the remaining 5 were negative for either *set1A* or *set1B*. However, all *S. flexneri* 1b and 1c strains were *set* negative (Fig 2, 3).

**Fig 2.**
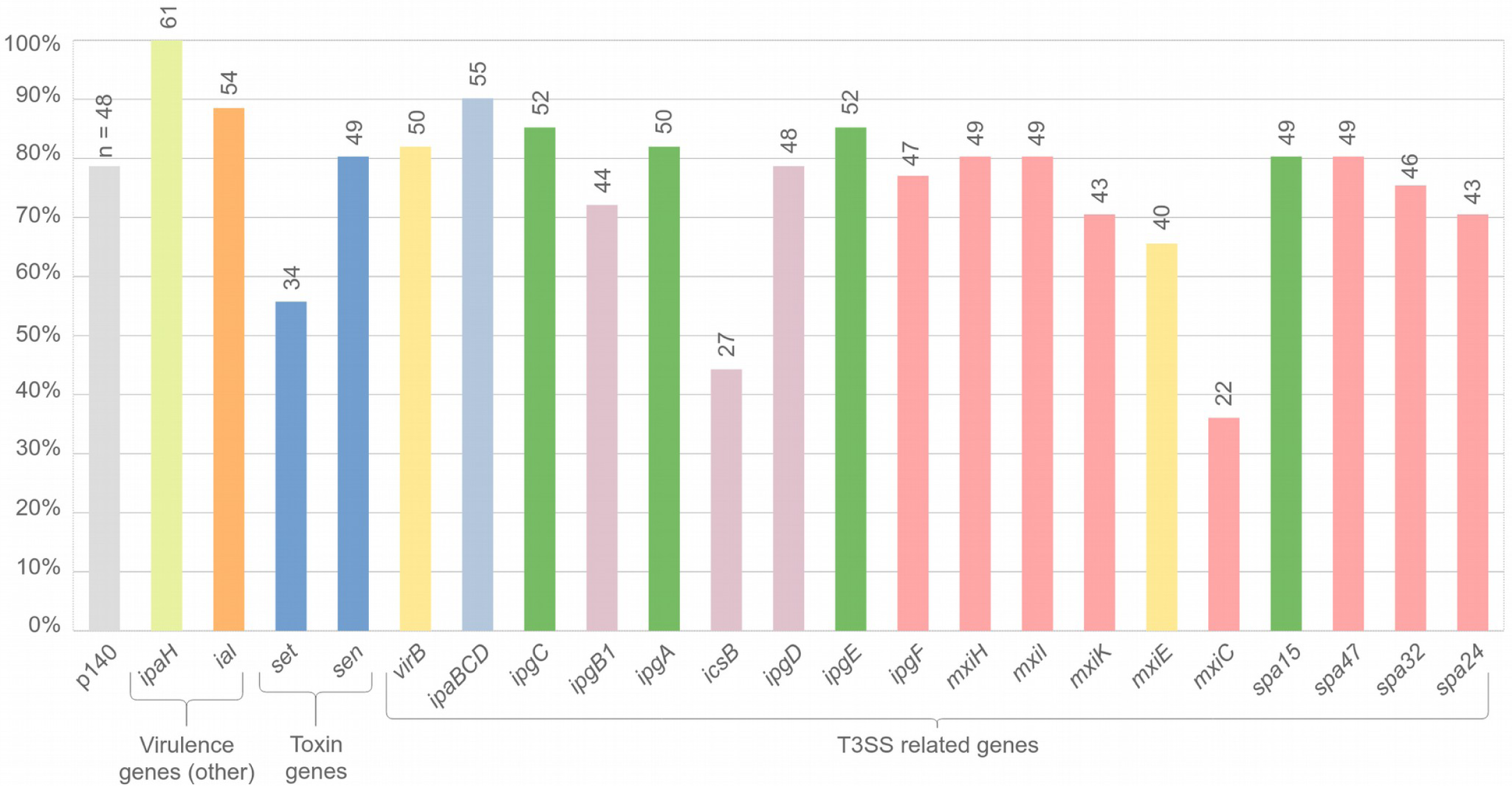
Prevalence p140, virulence, toxin and T3SS related genes in *S. flexneri* isolates. Positive results found in, p140 (n=48), *ipaH* (61/61), *ial* (54/61), *set* (34/61), *sen* (49/61), *virB* (50/61), *ipaBCD* (55/61), *ipgC* (52/61), *ipgB1* (44/61), *ipgA* (50/61), *icsB* (27/61), *ipgD* (48/61), *ipgE* (52/61), *ipgF* (47/61), *mxiH* (49/61), *mxiI* (49/61), *mxiK* (43/61), *mxiE* (40/61), *mxiC* (22/61), *spa15* (49/61), *spa47* (49/61), *spa32* (46/61), *spa24* (43/61).

**Fig 3.**
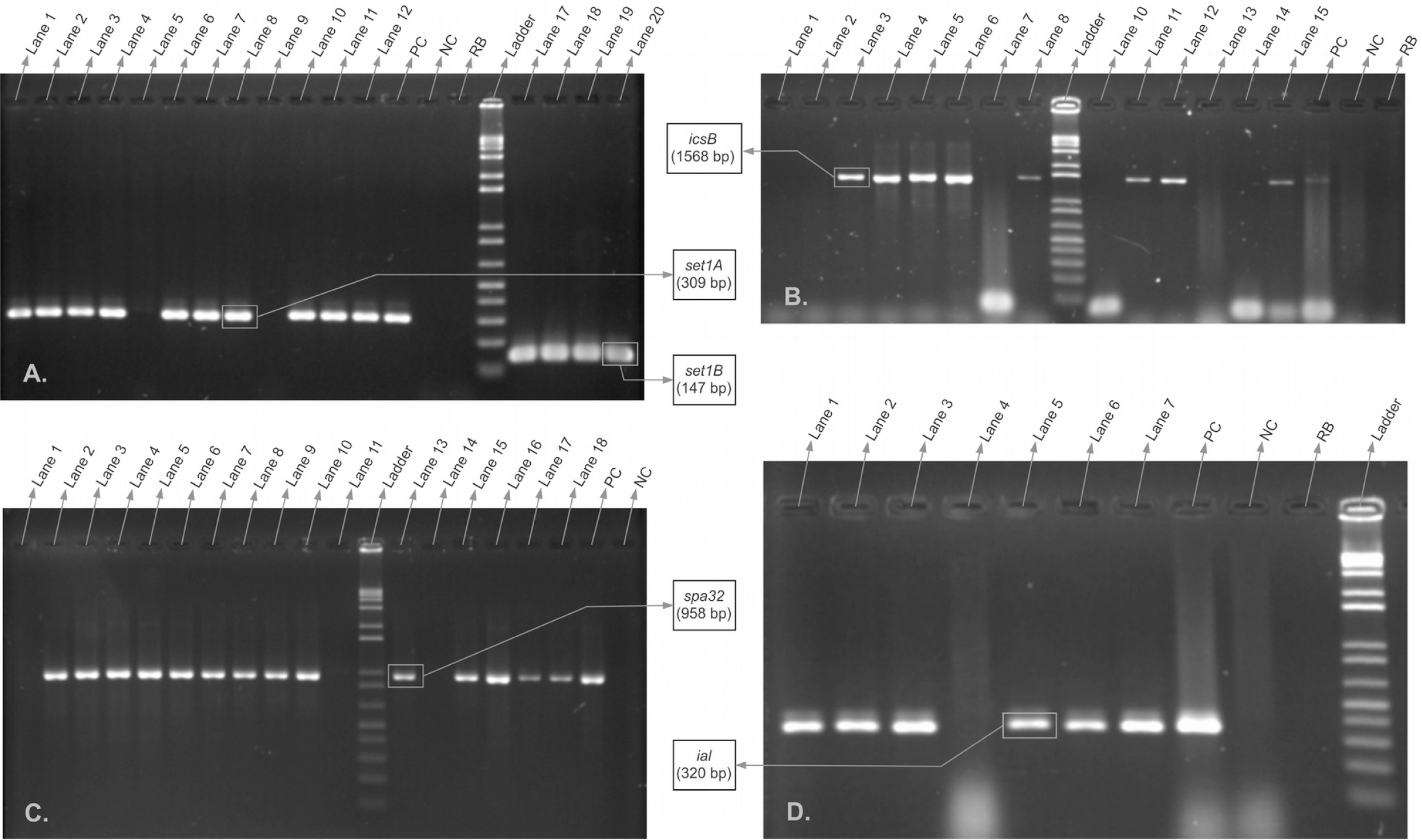
Gel electrophoresis of PCR products of representative *S. flexneri* strains. Agarose gel electrophoresis of (A) *set1A, set1B* genes: Lane 1 – K1063 – *S. flexneri* 2a, Lane 2 – K1057 – *S. flexneri* 2a, Lane 3 – K1053 – *S. flexneri* 2b, Lane 4 – K1044 – *S. flexneri* 2b, Lane 5 – K842 – *S. flexneri* 1c, Lane 6 – K662 – *S. flexneri* 2b, Lane 7 – K658 – *S. flexneri* 2b, Lane 8 – K645 – *S. flexneri* 2b, Lane 9 – K632 – *S. flexneri* 4, Lane 10 – K629 – *S. flexneri* 2b, Lane 11 – K583 – *S. flexneri* 2b, Lane 12 – K570 – *S. flexneri* 2b, Lane 13 – Positive control (PC), Lane 14 – Neative control (NC), Lane 15 – Reagent blank (RB), Lane 16 – 1 kb plus DNA ladder, Lane 17 – K1063 – *S. flexneri* 2a, Lane 18 – K1057 – *S. flexneri* 2a, Lane 19 – K1053 – *S. flexneri* 2b, Lane 20 – K1044 – *S. flexneri* 2b; (B) *icsB* gene: Lane 1 – K1044 – *S. flexneri* 2b, Lane 2 – K842 – *S. flexneri* 1c, Lane 3 – K662 – *S. flexneri* 2b, Lane 4 – K658 – *S. flexneri* 2b, Lane 5 – K649 – *S. flexneri* 2a, Lane 6 – K645 – *S. flexneri* 2b, Lane 7 – K632 – *S. flexneri* 4, Lane 8 – K629 – *S. flexneri* 2b, Lane 9 – 1 kb plus DNA ladder, Lane 10 – K583 – *S. flexneri* 2b, Lane 11 – K570 – *S. flexneri* 2b, Lane 12 – K569 – *S. flexneri* 2b, Lane 13 – K553 – *S. flexneri* 4, Lane 14 – K425 – *S. flexneri* 2b, Lane 15 – K270 – *S. flexneri* 3a, Lane 16 – Positive control, Lane 17 – Negative control, Lane 18 – Reagent blank; (C) *spa32* gene: Lane 1 – K842 – *S. flexneri* 1c, Lane 2 – K662 – *S. flexneri* 2b, Lane 3 – K658 – *S. flexneri* 2b, Lane 4 – K649 – *S. flexneri* 2a, Lane 5 – K645 – *S. flexneri* 2b, Lane 6 – K632 – *S. flexneri* 4, Lane 7 – K629 – *S. flexneri* 2b, Lane 8 – K583 – *S. flexneri* 2b, Lane 9 – K570 – *S. flexneri* 2b, Lane 10 – K569 – *S. flexneri* 2b, Lane 11 – K553 – *S. flexneri* 4, Lane 12 – 1 kb plus ladder, Lane 13 – K425 – *S. flexneri* 2b, Lane 14 – K270 – *S. flexneri* 3a, Lane 15 – K151 – *S. flexneri* 2b, Lane 16 – K102 – *S. flexneri* 2b, Lane 17 – K53 – *S. flexneri* 1c, Lane 18 – K6 – *S. flexneri* 3b, Lane 19 – Positive control, Lane 20 – Negative control; (D) *ial* gene: Lane 1 – K629 – *S. flexneri* 2b, Lane 2 – K583 – *S. flexneri* 2b, Lane 3 – K570 – *S. flexneri* 2b, Lane 4 – K553 – *S. flexneri* 4, Lane 5 – K270 – *S. flexneri* 3a, Lane 6 – K151 – *S. flexneri* 2b, Lane 7 – K53 – *S. flexneri* 1c, Lane 8 – Positive control, Lane 9 – Negative control, Lane 10 – Reagent blank, Lane 11 – 1 kb plus ladder. The expected positions of the PCR product of genes are shown with the indicators. In each case, 1 kb plus DNA ladder was used. As positive and negative controls, YSH6000 *S. flexneri* 2a, and *E. coli* ATCC 25922 strains were used, respectively.

### Assessment of T3SS related virulence genes

Of the 32 genes responsible for T3SS in *Shigella* species 18 different virulence genes - *virB, ipaBCD, ipgC, ipgB1, ipgA, icsB, ipgD, ipgE, ipgF, mxiH, mxiI, mxiK, mxiE, mxiC, spa15, spa47, spa32* and *spa24* - were analyzed in this study by PCR (Fig 2, 3). In these 61 strains, *ipaBCD* was found predominant and observed in 90% cases. The prevalence of *ipgC* and *ipgE* was 85% (52/61), *virB* and *ipgA* was 82% (50/61), *mxiK* and *spa24* was 70% (43/61), and for *mxiH, mxiI, spa15, spa47* the prevalence was 80% (49/61). The prevalence of the rest of the genes - *ipgD, ipgF, spa32, ipgB1, mxiE, icsB*, and *mxiC* - was 79, 77, 75, 72, 66, 44 and 36 percent respectively. Although 13 strains were found to be p140 negative, none of those strains were completely free of T3SS genes. At least one T3SS related gene was present in every one of them. In two strains - K-1080 (*S. flexneri* 2a) and K-842 (*S. flexneri* 1c) - this count was 17 and 6, respectively (Table 1).

**Table 1.**
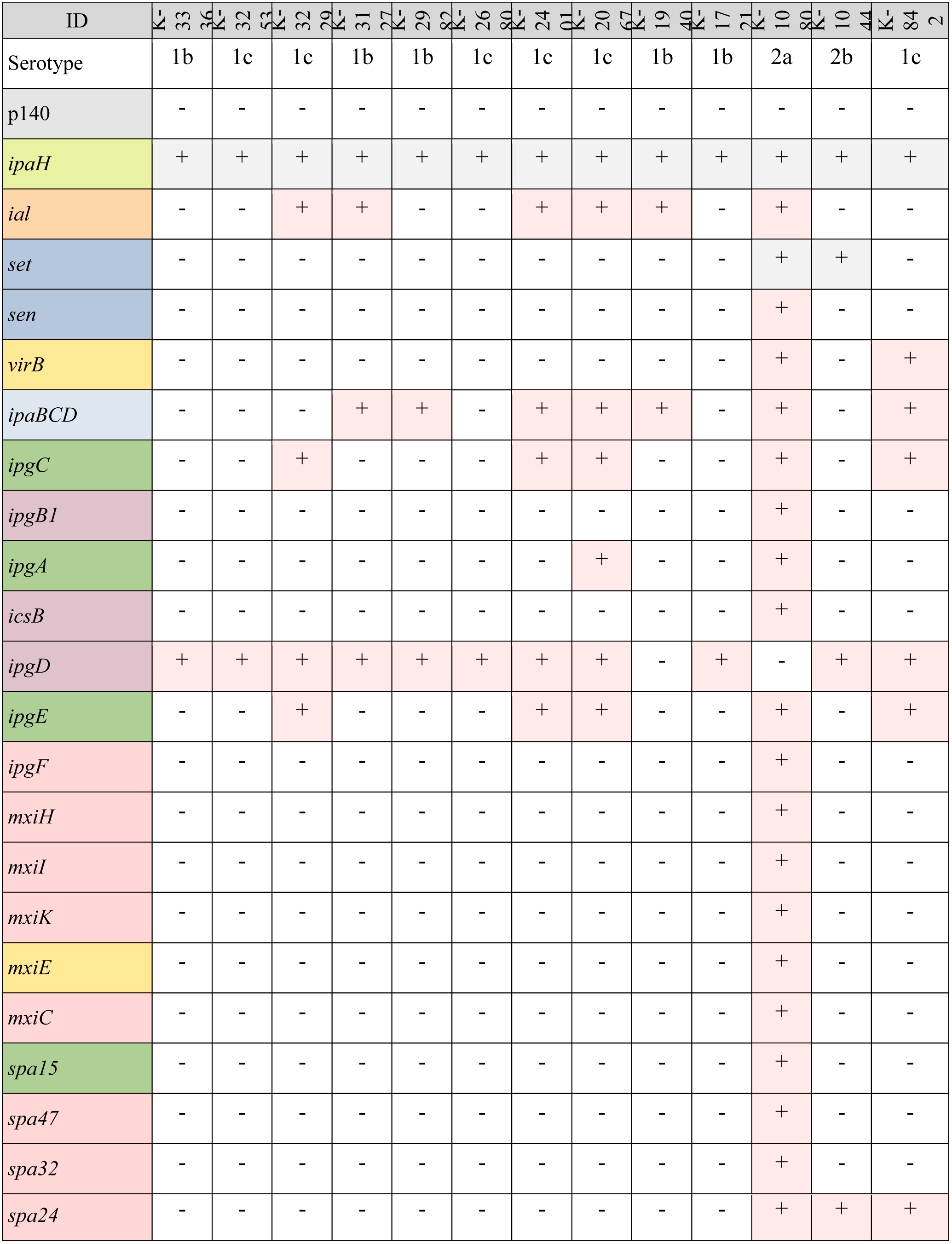
Status of tested genes in p140 negative *S. flexneri* strains.

### Clinical information

Relevant clinical information was obtained from the study database containing the history of present illness and clinical features observed in corresponding patients after admission, including abdominal pain (85.2%, 52/61), blood in stool (80.3%, 49/61), mucoid stool (80.3%, 49/61), fever (77%, 47/61), rectal straining (73.8%, 45/61), vomiting (45.9%, 28/61), presence of dehydration (21.3%, 13/61), sunken eyes (11.5%, 7/61), dry mouth (9.8%, 6/61), history of convulsion (3.2%, 2/61), loss of skin turgor (1.6%, 1/61), restlessness or irritability (1.6%, 1/61), and an overall increased incidence of diarrhoeal disease severity in 90.2% (55/61) cases. However, the status of the corresponding clinical features observed in the p140 negative strains are shown in Table 2.

**Table 2.** Status of clinical features in corresponding p140 negative strains.

| ID |  | 333 | 325 | 322 | 312 | 298 | 268 | 240 | 206 | 194 | 172 | 108 | 104 | K-842 |
| --- | --- | --- | --- | --- | --- | --- | --- | --- | --- | --- | --- | --- | --- | --- |
| Clinical features (frequencies) | Increased disease severity (11/13) | yes | no | yes | yes | no | yes | yes | yes | yes | yes | yes | yes | yes |
|  | Abdominal pain (9/13) | no | no | NA | yes | no | yes | yes | yes | yes | yes | yes | yes | yes |
|  | Blood in stool (10/13) | yes | no | no | yes | no | yes | yes | yes | yes | yes | yes | yes | yes |
|  | Mucoid stool (10/13) | yes | no | no | yes | no | yes | yes | yes | yes | yes | yes | yes | yes |
|  | Fever (8/13) | yes | no | yes | no | yes | yes | no | yes | yes | no | no | yes | yes |
|  | Rectal straining (8/13) | no | no | no | no | no | yes | yes | yes | yes | yes | yes | yes | yes |
|  | Vomiting (8/13) | yes | yes | yes | no | no | no | yes | no | yes | no | yes | yes | yes |
|  | Dehydration (2/13) | no | no | yes | no | no | no | no | no | no | no | no | no | yes |
|  | Sunken eyes (1/13) | no | no | yes | no | no | no | no | no | no | no | no | no | no |
|  | Dry mouth (1/13) | no | no | yes | no | no | no | no | no | no | no | no | no | no |
|  | Convulsion (0/13) | no | no | no | no | no | no | no | no | no | no | no | no | no |
|  | Loss of skin turgor (0/13) | no | no | no | no | no | no | no | no | no | no | no | no | no |
|  | Restlessness/irritability (0/13) | no | no | no | no | no | no | no | no | no | no | no | no | no |

**Table 3.** Summary of Chi-Square of independence.

| Gene | Clinical characteristics | Chi-Square test |  |  |  |
| --- | --- | --- | --- | --- | --- |
|  |  | Chi-square value | df | p-value |  |
|  |  |  |  | Asymptotic | Fisher's Exact |
| *ial* | Abdominal pain | 7.295<sup>a</sup> | 1 | - | 0.030 |
| *set* | Mucoid stool | 7.672<sup>b</sup> | 1 | - | 0.008 |
|  | Blood in stool | 9.243<sup>c</sup> | 1 | 0.002 | - |
|  | Rectal straining | 8.306<sup>d</sup> | 1 | 0.004 | - |
|  | Fever | 5.435<sup>f</sup> | 1 | 0.020 | - |
| *sen* | Mucoid stool | 7.672<sup>g</sup> | 1 | - | 0.008 |
|  | Blood in stool | 9.243<sup>h</sup> | 1 | 0.002 | - |
|  | Rectal straining | 8.306<sup>i</sup> | 1 | 0.004 | - |
|  | Fever | 5.435<sup>k</sup> | 1 | 0.020 | - |

a. 1 cell (25.0%) had expected count less than 5. The minimum expected count was 83. b. 1 cell (25.0%) had an expected count of less than 5. The minimum expected count was 4.87. c. 0 cells (0.0%) had an expected count of less than 5. The minimum expected count was 5.31. d. 0 cells (0.0%) had an expected count of less than 5. The minimum expected count was 7.08. f. 0 cells (0.0%) had an expected count of less than 5. The minimum expected count was 6.20. g. 1 cell (25.0%) had an expected count of less than 5. The minimum expected count was 4.87. h. 0 cells (0.0%) reported an expected count less than 5. The minimum expected count was 5.31. i. 0 cells (0.0%) had an expected count of less than 5. The minimum expected count was 7.08. k. 0 cells (0.0%) had an expected count of less than 5. The minimum expected count was 6.20. The minimum expected count was 6 with 20 df – degrees of freedom.

### Statistical analysis

Chi-Square test of independence was done to determine the association between the investigated genes and the clinical features observed in corresponding patients. The result was considered as statistically significant if the asymptotic significance (2-sided) was p<0.05. However, in the case of the expected cell value of <5, Fisher’s Exact test was considered (Table 2). In total, nine statistically significant associations were found (all p<0.05). Both of the enterotoxin genes - *set* and *sen* - were found highly associated with the following clinical features: mucoid stool (p=0.008), blood in stool (p=0.002) and rectal straining (p=0.004). These two toxin genes were also found to be associated with fever (p=0.020). And lastly, another association was found between the *ial* gene and abdominal pain (p=0.030).

## Discussion

In this study, 61 randomly selected strains of *S. flexneri* were analyzed. Plasmid profiling was done for all of them to detect the status of 140 MDa large virulence plasmid (p140). PCR assay was done individually to determine the status of virulence (*ipaH, ial*), toxin (*set, sen*) and T3SS related genes (*virB, ipaBCD, ipgC, ipgB1, ipgA, icsB, ipgD, ipgE, ipgF, mxiH, mxiI, mxiK, mxiE, mxiC, spa15, spa47, spa32, spa24*). In this study, we designed the primers for *mxiE* - a T3SS regulator - and *spa15* - a T3SS chaperone - genes with a web-based bioinformatics tool, primer3plus. The validity of these primers was confirmed, when the predicted product size correlated with the band size of the PCR product during gel electrophoresis analysis.

In these 61 strains, *ipaBCD* was observed in 90% cases. The prevalence of *ipgC* and *ipgE* was 85% (52/61), *virB* and *ipgA* was 82% (50/61), *mxiK* and *spa24* was 70% (43/61). For *mxiH, mxiI, spa15, spa47* this proportion was 80% (49/61). The prevalence of *ipgD, ipgF, spa32, ipgB1, mxiE, icsB* and *mxiC* was 79, 77, 75, 72, 66, 44 and 36 percent, respectively. *ipaH* was found in all strains, however the presence of the rest of the genes were variable. Out of the 48 p140 positive strains, the frequency of *ial, set, sen, virB, ipaBCD, ipgC, ipgB1, ipgA, icsB, ipgD, ipgE, ipgF, mxiH, mxiI, mxiK, mxiE, mxiC, spa15, spa47, spa32* and *spa24* were, 48, 32, 32, 48, 48, 47, 43, 48, 26, 37, 47, 46, 48, 48, 42, 39, 21, 48, 48, 45 and 40, respectively.

Apart from the increased incidence of diarrheal disease severity (90.2%, n=55), the clinical features observed in corresponding patients were: abdominal pain, blood in stool, mucoid stool, fever, rectal straining, vomiting, dehydration, sunken eyes, dry mouth, convulsion, loss of skin turgor, restlessness or irritability, with a prevalence of 85.2% (52/61), 80.3% (49/61), 80.3% (49/61), 77% (47/61), 73.8% (45/61), 45.9% (28/61), 21.3% (13/61), 11.5% (7/61), 9.8% (6/61), 3.2% (2/61), 1.6% (1/61) and 1.6% (1/61), respectively.

Theoretically, the T3SS related genes are located inside the 31 kb “entry region” of p140 (7)□, but none of these p140 negative strains were found devoid of these genes, completely (Table 1). One to five T3SS related genes were found invariably, while in K-842 (*S. flexneri* 1c) and K-1080 (*S. flexneri* 2a), 6 and 17 of such T3SS related genes were detected, respectively. Similarly, multiple clinical features relevant to shigellosis were observed in these p140 negative strains at a varying degree (Table 2). And, in K-1080 (*S. flexneri* 2a) and K-842 (*S. flexneri* 1c), the clinical traits analyzed in this study were found at a higher proportion, including abdominal pain, bloody mucoid stool, fever, rectal straining, vomiting and an increased state of diarrheal disease severity. As these strains were pathogenic even though they were missing p140, we suspect that these T3SS related genes might be integrated elsewhere within the bacterial genome, which could be mediated by transposons or insertion sequences (25)□.

To determine any significant association between any particular clinical feature and tested genes, the Chi-Square test of independence was done and several statistically significant (p<0.05) associations were found. Both of the *Shigella* enterotoxin genes - *set* and *sen* - were found highly significantly associated with the presence of blood in stool (p=0.002), mucoid stool consistency (p=0.008) and rectal straining (p=0.004). However, they were also found significantly associated with fever (p=0.02). The chromosomal gene *set* is composed of two domains - *set1A* and *set1B* - encoding ShET1A and ShET1B proteins, while ShET1A is responsible for the secretory activity, and ShET1B causes the irreversible binding of the toxin to the enterocyte receptors (26)□. In contrast, the *sen* gene is plasmid-borne which codes for ShET2 (*Shigella* enterotoxin 2), causing epithelial inflammation by contributing to the release of Interleukin-8 (IL-8) from the gut epithelium (27)□. As IL-8 is a chemokine, it attracts and activates the neutrophil at the inflammatory region, causing injury to the nearby cells, such as rectal epithelium (28)□. This chain of events is probably responsible for the clinical features that we observed in this study, such as bloody mucoid stool, rectal straining, and fever, which also corroborates with the results of the previous studies.

The *ial* gene has been reported to direct the epithelial cell penetration by the organism (29)□, and in our study, we also found *ial* to be significantly associated with abdominal pain (p=0.03), which reinforces the findings observed in past studies.

Nevertheless, in several studies, shigellosis has been reported to be associated with convulsion (30)□, but we found no significant statistical correlation in this study. To verify this matter a larger sample size with additional clinical data is required.

## Conclusion

In our study, the two *Shigella* enterotoxin genes - *set* and *sen* - along with another virulence gene - *ial* - were found significantly associated with diverse clinical features relevant to shigellosis, including bloody-mucoid stool, rectal straining, fever, and abdominal pain. In the future, further studies with a larger sample size are needed to decipher how a p140 negative *Shigella* strain can develop shigellosis.

## Acknowledgments

We gratefully acknowledge the core donors of icddr,b – the Government of the People’s Republic of Bangladesh, Global Affairs Canada (GAC), Swedish International Development Cooperation Agency (Sida) and the Department for International Development (UK Aid) – for providing unrestricted support to icddr,b for its operations and research.

